# The Interictal Suppression Hypothesis in Focal Epilepsy: Electrographic and Structural Evaluation

**DOI:** 10.1101/2022.06.27.497765

**Authors:** Graham W. Johnson, Derek J. Doss, Victoria L. Morgan, Jared S. Shless, Danika L. Paulo, Hakmook Kang, Sarah K. Bick, Shawniqua Williams Roberson, Mark T. Wallace, Dario J. Englot

**Affiliations:** Vanderbilt University Department of Biomedical Engineering - Nashville, Tennessee, 37235 USA; Vanderbilt University Institute of Imaging Science (VUIIS) - Nashville, Tennessee, 37232 USA; Vanderbilt Institute for Surgery and Engineering (VISE) – Nashville, Tennessee, 37235 USA; Vanderbilt University Medical Center Department of Neurological Surgery - Nashville, Tennessee, 37232 USA; Vanderbilt University Department of Electrical Engineering and Computer Science - Nashville, Tennessee, 37235 USA; Vanderbilt University Medical Center Department of Neurology - Nashville, Tennessee, 37232 USA; Vanderbilt University Medical Center Department of Radiological Sciences - Nashville, Tennessee, 37232 USA; Vanderbilt University Department of Biostatistics - Nashville, Tennessee, 37232 USA; Vanderbilt University Department of Hearing and Speech Sciences - Nashville, Tennessee, 37232 USA; Vanderbilt University Department of Psychology - Nashville, Tennessee, 37232 USA; Vanderbilt University Department of Psychiatry and Behavioral Sciences - Nashville, Tennessee, 37232 USA; Vanderbilt University Department of Pharmacology - Nashville, Tennessee, 37232 USA

## Abstract

Why are people with focal epilepsy not constantly seizing? Previous molecular work has implicated gamma-aminobutyric acid balance as integral to seizure generation and termination, but is the high-level distributed brain network involved in suppressing seizures? Recent intracranial electrographic evidence has suggested that seizure onset zones have an increased inward connectivity. Accordingly, we hypothesize that seizure onset zones are actively suppressed by the rest of the brain network during interictal states.

We tested this hypothesis on 81 subjects with drug resistant focal epilepsy undergoing presurgical evaluation. We utilized intracranial electrographic resting-state and neurostimulation recordings to evaluate the network connectivity of seizure onset, propagative, and non-involved regions. We then utilized diffusion imaging to acquire estimates of white matter connectivity to evaluate structure-function coupling effects on connectivity findings. Finally, using our observations, we generated a resting-state classification model to assist clinicians in detecting seizure onset and propagative zones without the need for multiple ictal recordings.

Our findings indicate that seizure onset and propagative zones demonstrate markedly increased inward connectivity and decreased outward connectivity on both resting-state and neurostimulation analyses. When controlling for distance between regions, the difference between inward vs. outward connectivity remained stable up to 80 mm between brain connections. Structure-function coupling analyses revealed that seizure onset zones exhibit abnormally enhanced coupling (hypercoupling) of surrounding regions compared to presumably healthy tissue. Using these observations, our classification models achieved a maximum held-out testing set accuracy of 92.0±2.2%.

These results indicate that seizure onset zones are actively segregated and suppressed by a widespread brain network. Furthermore, this electrographically observed functional suppression is disproportionate to any observed structural connectivity alterations of the seizure onset zones. These findings have implications for the identification of seizure onset zones using only brief resting-sate recordings to reduce patient morbidity and augment the presurgical evaluation of drug resistant epilepsy. Furthermore, testing of the interictal suppression hypothesis can provide insight into potential new resective, ablative, and neuromodulation approaches to improve surgical success rates in those suffering from drug resistant focal epilepsy.

## Introduction

Drug-resistant focal epilepsy (DRE) accounts for 30-40% of the estimated 50 million cases of epilepsy worldwide. ^1, 2^ If medication fails to control seizures, patients may elect to pursue surgical treatments including resection,^3, 4^ ablation,^5, 6^ or neurostimulation.^7, 8^ Traditional presurgical evaluation includes identification of seizure-onset zones (SOZs) using seizure semiology, non-invasive neuroimaging and scalp EEG.^9^ SOZs are defined as observed sites of ictal onset and are candidates for an electro-clinically defined “epileptogenic zone” that could theoretically be removed to render the patient seizure free.^10^ If these techniques fail to accurately localize SOZs, then invasive monitoring with stereotactic-electroencephalography (SEEG) can be pursued.^11^ To properly localize SOZs with SEEG, the patient must remain in the hospital for days to weeks to record many seizures (ictal events).^12, 13^ To reduce the morbidity of long intracranial recordings, brief resting-state (interictal) SEEG analysis has shown promise in identifying SOZs.^14–20^

Resting-state SEEG studies often utilize the conceptual framework of epilepsy as a disorder of interconnected brain nodes (i.e. a network).^21–26^ Specifically, recent work has shown that SOZs exhibit increased interictal *inward* connectivity from other nodes of the brain.^20, 27^ This work has led to what we outline as the Interictal Suppression Hypothesis (ISH), which states that the SOZ is tonically supressed by other areas of the brain during resting-state to prevent seizure activity. The premise of this hypothesis is intuitive based off the clinical observation that people with DRE are not continuously seizing - perhaps there is a an interictal widespread functional organization of the brain that is actively suppressing epileptiform activity. Further, neurophysiological evidence for this hypothesis has been outlined in animal models showing GABA (γ-aminobutyric acid) mediated receptor tonic interictal inhibition of SOZs.^28, 29^ In humans, we can test the ISH by using SEEG electrodes to investigate the connectivity of the epileptic network. SEEG signal observations are electrical phenomena of underlying physiological activity and provide direct sampling of brain tissue to provide insights into the high-level functional organization of the brain in humans living with DRE.

Beyond resting-state recordings, SEEG electrodes can also be used to apply electrical stimulation to the brain. Specifically, single-pulse electrical stimulation (SPES) can be used to simultaneously stimulate one region and record the post-stimulation EEG signal at all other implanted regions.^30–34^ SPES provides another paradigm to test the ISH. Importantly, not only can directionality be inferred from SPES, but relative elevation or attenuation of oscillatory activity can also be quantified to gain insight into the relative excitatory and inhibitory nature of functional connections.

Finally, the spatial and anatomical characteristics of the network have been shown to impact the resting-state and stimulation-derived network characteristics. Specifically, past work has shown short-range connections to be driving increased SOZ connectivity,^19^ and diffusion-weighted imaging (DWI) derived structural connectivity helps explain the network’s response to SPES.^35, 36^ Using these techniques, we can evaluate the effects of distance and structural connectivity to further characterize the functional epileptic network.

Overall, this work aims to test the ISH using resting-state and stimulation-based electrophysiology, controlling for spatial and anatomical variability. Better characterization of both SOZs using only interictal data may reduce patient morbidity by reducing reliance on ictal recordings and provide insight into the biophysical pathophysiology of the epileptic network.

## Materials and methods

### Participants & Seizure Network Designations

We included 81 patients with DRE undergoing SEEG presurgical evaluation at Vanderbilt University Medical Center (VUMC). This study was approved by Vanderbilt’s Institutional Review Board and all patients underwent informed consented. The diagnosis of DRE and decision to pursue SEEG was determined by our medical center’s standard multidisciplinary process, including epileptologists, neurosurgeons, and neuropsychologists. This process included analyzing patient history, seizure semiology, video EEG, MRI, positron emission tomography, memory/language localization by functional MRI or Wada, and neuropsychological testing. Electrode (Ad-Tech, or PMT Cooperation) trajectories were planned by treating physicians according to standard clinical care at VUMC using CRAnialVault Explorer (CRAVE; Vanderbilt University, Nashville, Tennessee).^37^

To assign seizure network designations to the SEEG contacts, the treating epileptologist reviewed all electroclinical ictal events during the SEEG monitoring. Seizure onset zones (SOZs) were defined as SEEG contact(s) with the first electrographic epileptiform changes observed for a clinically significant ictal event. Early propagation zones (PZs) were defined as SEEG contacts with electrographic epileptiform activity spread within 10 seconds of ictal onset. Finally, non-involved (Non) SEEG contacts were defined as belonging to neither SOZ or PZ designations.^19, 20^ Finally, to quantify response to surgical intervention we calculated the Engel Surgical Outcome Scale for subjects who received a resective or ablative surgery and had follow-up data at least one year following surgery.^38^ Patients who received a neuromodulatory treatment were excluded from analyses pertaining to surgical outcome.

### Resting-state SEEG Connectivity

We calculated undirected and directed SEEG resting-state connectivity for all 81 subjects as outlined in our previous works.^19, 20^ Briefly, we collected 20 minutes of resting-state data on SEEG post-implantation day one or two with the subject lying flat with their eyes closed trying not to fall asleep. We then filtered the data with a 1-59, 61-80 Hz bandpass Butterworth filter using Matlab’s filtfilt function (MathWorks inc., Natick, MA, USA) to remove direct current offset and power line interference. The signals were referenced using a bipolar montage. For simplicity, each bipolar pair referenced signal will henceforth be referred to as ‘node’, and the connectivity between two nodes will be referred to as an ‘edge’. To reduce potential segmentation bias, we did not use a brain atlas for any portion of these analyses.^39^ We then segmented the 20-minute resting-state epoch into 10 two-minute epochs. All metric calculations described below were averaged across these 10 two-minute epochs.

To assess undirected resting-state SEEG connectivity, we calculated imaginary coherence (ImCoh) for all edges.^19^ For directed connectivity analysis, we calculated partial directed coherence (PDC) for all edges.^20^ Specifically, a node’s undirected connectivity was defined as the average ImCoh value of that node’s edges to all other nodes. A node’s outward connectivity was defined as the average outward PDC of edges to all other nodes. Correspondingly, a node’s inward connectivity was defined as the average inward PDC of edges from all other nodes. To quantify the overall direction of a node’s connectivity, reciprocal connectivity was calculated as the subtraction of inward minus outward connectivity for each node. All connectivity was analyzed in the theta [4-7 Hz], alpha [8-12 Hz], beta [12-30 Hz], and gamma [31-80 Hz] frequency bands.

Additionally, to evaluate the effect of distance on connectivity measurements, we calculated the Euclidean distance between all nodes (centroid of the bipolar pair) using the CRAVE software. We then re-calculated all functional connectivity measurements with Euclidean edge distance thresholds of 5-20, 20-35, 35-50, 50-65, 65-80 mm. These distance thresholds were chosen to include a roughly equal number of SEEG nodes across a wide range of Euclidean distances for the entire subject cohort. In this manner, we were able to observe the effect of distance between the SEEG nodes on the functional connectivity measurements.

### Single-Pulse Electrical Stimulation (SPES) Connectivity

To further test the ISH, we collected neurostimulation data from the most recently enrolled subset of 23 patients in the cohort. Specifically, we conducted SPES with every SEEG node in gray matter for each patient. We used a 10 second train of 1 Hz, 300 microsecond, biphasic pulses at 3.0 milliamps with a recording sampling rate of 512 Hz. We filtered raw SEEG data using Matlab’s filtfilt function with Butterworth filter passbands of 1-59, 61-80 Hz. We then parsed the data into epochs of 5-305 ms following each stimulation pulse. This post-stimulation epoch has been previously outlined to capture the majority of SPES induced electrographic changes.^31^ The initial 5 ms was omitted to avoid stimulation artifact.^40^

We then created SPES-derived connectivity matrices by computing the power spectral density (PSD) for the frequency bands outlined in “Resting-State SEEG Connectivity”. The PSD for each node was normalized to that node’s pre-stimulation baseline. The outward connectivity for a given node was assigned as the PSD change from baseline within a given frequency range for all non-stimulated SEEG nodes when that node was stimulated. Conversely, the inward connectivity for a given node was assigned as the PSD change from pre-stimulation baseline in the frequency band of interest for that node when all other nodes had been stimulated separately. All PSD measurements were averaged across the 10 redundant stimulation pulses to increase the signal to noise ratio. We did not attempt to analyze any frequencies below 4 Hz due to the short sampling window post-stimulation.

### Diffusion Imaging Collection & Preprocessing

We collected DWI data using high angular resolution diffusion imaging (HARDI) acquisition with 92 b-vector directions at a b-value of 1600 s/mm^2^. We preprocessed the DWI data with a pipeline built around MRTrix3^41^, FSL^42^, and ANTs^43^ software packages.^44^ First, the diffusion data was denoised with the Marchenko-Pastur PCA method.^45–47^ The images were then intensity-normalized to the first image and concatenated for further processing. No reverse phase encoded images were acquired, but corresponding T1 images of the subjects were available. Thus, a T1 image was used to generate a synthetic susceptibility-corrected b0 volume using SYNB0-DISCO, a deep learning framework by Schilling et al.^48^ This synthetic b0 image was used in conjunction with FSL’s topup to correct for susceptibility-induced artifacts in the diffusion data. FSL’s eddy algorithm was then be used to correct for motion artifacts and eddy currents and to remove outlier slices.^49–52^

### Patient Specific SEEG-Based Structural Connectivity: The SWiNDL Technique

Next, we sought to evaluate the structure-function coupling between brain regions sampled by SEEG. To accurately quantify the structural connectivity between SEEG electrodes, we implemented a novel technique termed “<u>S</u>ubsampling <u>W</u>hole-Brain Tractography with <u>i</u>EEG <u>N</u>ear-Field <u>D</u>ynamic <u>L</u>ocalization” (SWiNDL).We developed this technique to obtain a patent-specific SEEG contact-to-contact structural connectome by utilizing the group analysis advantages of anatomically constrained whole-brain tractography and spherical deconvolution filtering of tractograms (SIFT2).^53^

First, we performed whole-brain tractography using MRtrix3 with SIFT2 weighting to generate ten million streamlines.^41, 53^ Briefly this included brain extraction from the T1 image,^54^ bias field correction,^55^ and registration to DWI^56^. The T1 image was then be used to create 5-tissue type (5TT)^57–59^ segmentations. Spherical deconvolution was performed on the preprocessed DWI to get direct fiber orientation density function for every voxel.^60, 61^ We then performed probabilistic tractography^62^ utilizing the 5TT volume to perform anatomically constrained tractography^63^ with dynamic seeding.^49^ The resulting tractogram was then corrected using spherical-deconvolution informed filtering of tractograms (SIFT2) for improved biological accuracy of white-matter representation.^53, 64^

We then localized each patient’s SEEG electrodes using CRAVE. We assigned a three-dimensional Gaussian probability distribution around each SEEG contact to model the volume of tissue electrically sampled by that electrode – we used a 95% attenuation of the probability at 12 mm based off previous electrophysiological work characterizing the spatial dynamics of local field potentials.^65^ For the connection between each pair of SEEG contacts (i.e. an edge), we then took the raw tractogram of 10 million streamlines and assigned dynamic weights [0-1] to each streamline based off the location of the endpoints relative to the SEEG contacts. For example, a streamline would be highly sampled with a weight close to 1.0 if each endpoint were near the center of a contact. Whereas, if one or both endpoints were far from the two SEEG contacts of interest, then that streamline would be assigned a weight close to 0.0 based off the minimum Gaussian distribution value. The weighted sum of the 10 million streamlines for each bivariate pair of contacts is summed to achieve the SWiNDL structural connectivity for that edge. SWiNDL allows for patient-specific SEEG contact-level connectomes that are dynamically weighted to the proximity of streamlines to the contacts and does not rely on inconsistent probabilistic tractography seeding between patients.

Using the SWSiNDL structural connectivity, we calculated the average connectivity for SOZs, PZs, and non-involved regions across a subset of 26 subjects that had available diffusion imaging. Finally, we evaluated the structure-function coupling by calculating the ratio of structural/functional connectivity for each SEEG edge within a patient. Well coupled structural and functional connections are expected to have a relative ratio of unity. Thus, we termed any connections that had disproportionality high functional connectivity compared to average or low structural connectivity as “hypercoupled”.

### SEEG Node Classification Model

Using the functional and structural connectivity metrics outlined above, we sought to test the efficacy of a support vector machine (SVM) classification of individual SEEG nodes (i.e. bipolar channel pairs) as SOZ, PZ or non-involved. For all SVM model development and evaluation we utilized a 5-fold nested cross validation scheme (**Figure 6A**) that utilizes a completely held out test set for each fold of model training and validation. This way, we can repeatedly test a model’ s generalizability while minimizing overfitting the training data. All fold splits were conducted at the patient-level.

First, we developed a model to classify each SEEG node as SOZ, PZ, or non-involved using only the resting-state functional connectivity metrics of ImCoh and PDC (6928 SEEG nodes for 81 subjects). Next, we sought to evaluate if the model improved with the addition of patient-specific structural connectivity metrics obtained from SWiNDL evaluation of DWI (2051 SEEG nodes from 26 subjects). Finally, we trained a model using only functional data from subjects who received resective or ablative therapy and achieved Engel I outcomes at least one year after surgery (1431 SEEG nodes from 19 subjects), and then evaluated the model on subjects who received resective or ablative therapy with Engel II-IV surgical outcomes (1265 SEEG nodes from 15 subjects).

## Results

### SOZs and PZs Exhibit Evidence of Suppression at Rest

The resting-state SEEG analysis with 81 subjects revealed that SOZs and PZs exhibit significantly increased undirected connectivity compared to non-involved regions. These results were consistent across all frequency bands. The alpha band results are depicted in **Figure 1A** (one-way ANOVA p-value=2.13e-3 with post-hoc multiple pairwise t-test comparisons depicted in figure). For the directed analysis, SOZs and PZs demonstrated markedly elevated inward connectivity and lower outward connectivity compared to non-involved regions - the mean PZ inward and outward connectivity was significantly intermediate to the SOZ and non-involved mean (**Figure 1B-C**, one-way ANOVA p-values=1.75e-12, 4.95e-10 for inward and outward respectively). The reciprocal connectivity (inward – outward) exhibited an increased significance with a one-way ANOVA p-value of 3.13e-13 (**Figure 1D**). As with undirected connectivity, the directed results were consistent across all frequency bands (one-way ANOVA p-value<1e-10 for all bands). These findings suggest that high-level functional organization of the epileptic network, as observed on resting-state SEEG, exhibits a very high functional suppression of SOZs and PZs characterized by elevated inward connectivity and lower outward connectivity.

**Figure 1:**
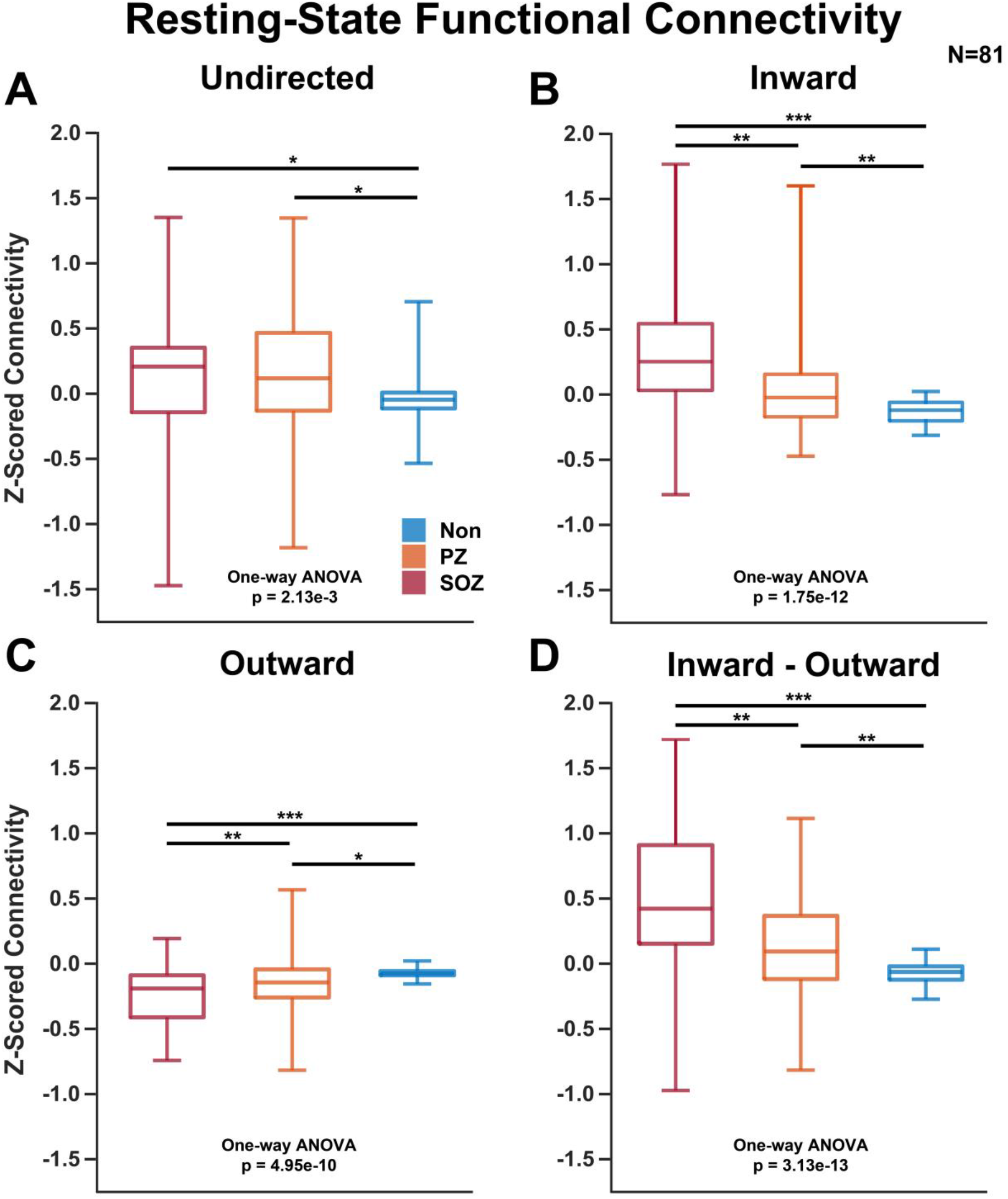
Resting-state SEEG connectivity. **A)** Undirected alpha-band imaginary coherence was elevated for SOZs and PZs (one-way ANOVA p-value=2.13e-3 with post-hoc multiple pairwise t-test comparisons significant for SOZ-Non and PZ-Non. **B)** Inward partial directed coherence strength was elevated significantly for SOZs and PZs (one-way ANOVA p-value=1.75e-12 with post-hoc multiple pairwise t-test comparisons significant between all three groups. **C)** Outward partial directed coherence strength was significantly lower for SOZs and PZs (one-way ANOVA p-value=4.95e-10 with post-hoc multiple pairwise t-test comparisons significant between all three groups. **D)** Inward – outward (reciprocal) connectivity exhibited a stronger signal to that of inward or outward separately (one-way ANOVA p-value=3.13-13 with post-hoc multiple pairwise t-test comparisons significant between all three groups). *p<5e-2, **p < 5e-3, ***p< 5e-6. N=81 subjects.

### Low Frequency Power is Attenuated in SOZs When the Network is Stimulated

When conducting SPES on non-SOZ SEEG contacts, we observed that the SOZ power was markedly attenuated compared to baseline and significantly attenuated compared to non-involved regions in the lowest frequency band measured (theta, 8-12 Hz) for the 23 patients consented for neurostimulation (**Figure 2A**, SOZ single-population t-test p-value=2.28e-4, one-way ANOVA p-value=2.50e-3, post-hoc pairwise t-test comparison SOZ-Non p-value=1.80e-3). The SOZ and PZ power was not significantly altered in the alpha band [8-12 Hz] (**Figure 2B**). The SOZ and PZ power was elevated compared to baseline in both beta [13-30 Hz] and gamma bands [31-80 Hz], but post-hoc multiple comparisons revealed that the SOZ elevation in power was only significantly different from non-involved regions in the beta band and only from PZs in the gamma band (**Figure 2C-D**, beta: SOZ single-population t-test p-value=9.00e-3, one-way ANOVA p-value=2.98e-2, post-hoc pairwise t-test comparison SOZ-

**Figure 2:**
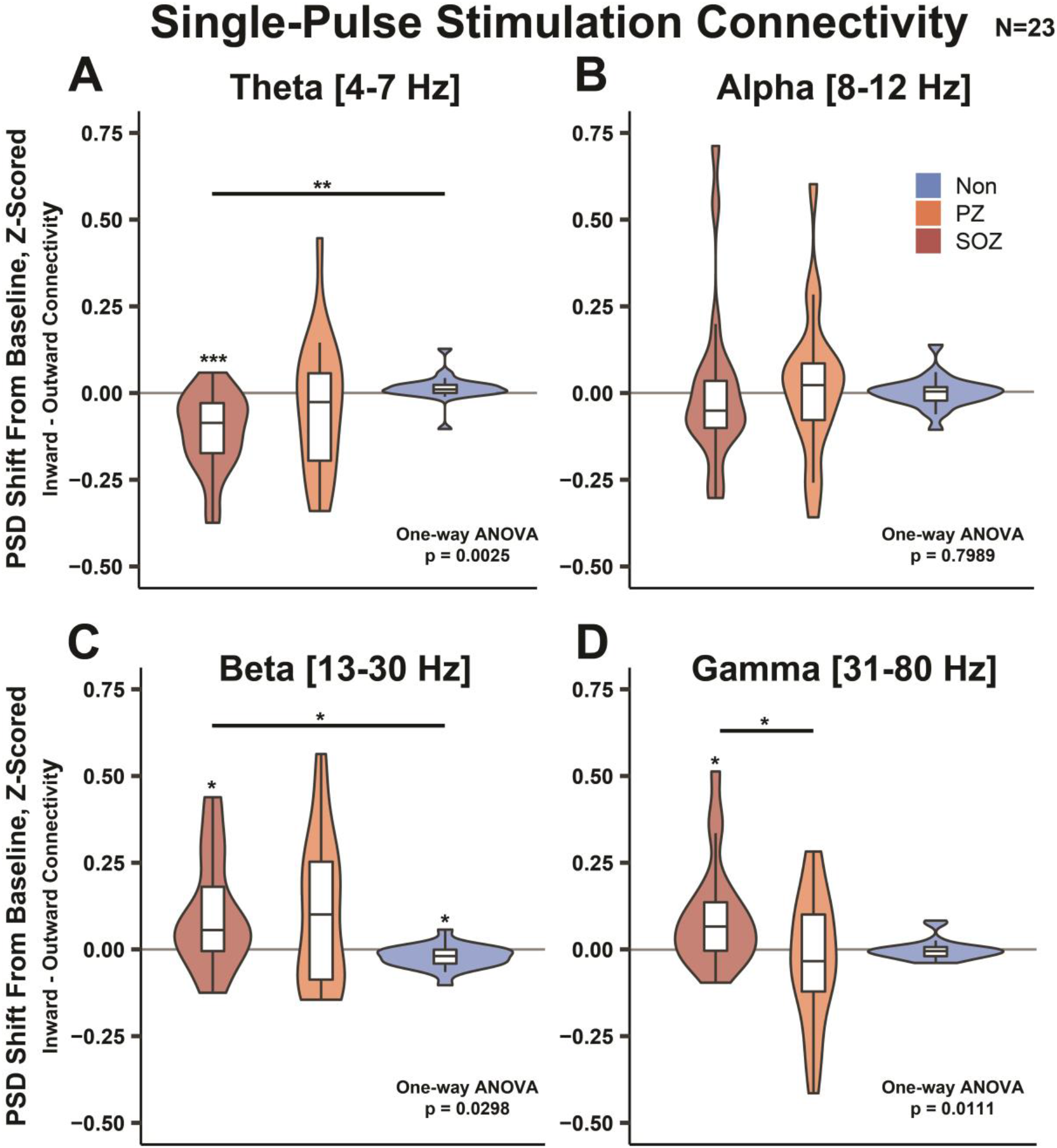
SPES connectivity. Frequency-band specific low-frequency stimulation induced change in power spectral density from pre-stimulation baseline. **A)** SOZ theta power is reduced when non-SOZ SEEG contacts are stimulated. **B)** Alpha band power was not observed to be significantly altered during stimulation. **C-D)** Beta and gamma band power in SOZs were elevated when non-SOZs were stimulated. *p<5e-2, **p < 5e-3, ***p< 5e-6. N=23 subjects.

Non p-value=2.24e-2; gamma: SOZ single-population t-test p-value=6.70e-3, one-way ANOVA p-value=1.11e-2, post-hoc pairwise t-test comparison SOZ-PZ p-value=1.33e-2). These results indicate that low-frequency power is markedly attenuated in SOZs, but not PZs or non-involved regions when other nodes of the network are stimulated.

### Inward vs. Outward Connectivity Remains Constant Over Long Distances

When evaluating the effects of distance on functional connectivity on all 81 patients in the cohort, we observed that both undirected and directed connectivity dropped significantly as Euclidean edge length increased (**Figure 3A-C**, two-way repeated measures ANOVA distance effect p-value<1e-10 for undirected inward and outward connectivity). Furthermore, the SOZ, PZ and non-involved region undirected connectivity 95% confidence intervals of the mean overlapped significantly for edge distances greater than 20 mm. However, inward and outward connectivity for SOZs, PZs and non-involved regions remained significantly different from each other over the span of Euclidean edge distances (two-way repeated measures ANOVA group effect p-value<1e-10 for inward and outward). Calculating the reciprocal connectivity over distance, we observed that the relationship between inward vs. outward connectivity remained consistent over the span of distances as can be observed by the flattening of the lines in **Figure 3D** (two-way repeated measures ANOVA group effect p-value=2.6e-12, interaction p-value=1.15e-6). These results indicate that the network’s potential functional suppression of SOZs and PZs, as measured by increased inward connectivity and decreased outward connectivity, scales proportionally as distance increases – thus suggesting whole network involvement.

**Figure 3:**
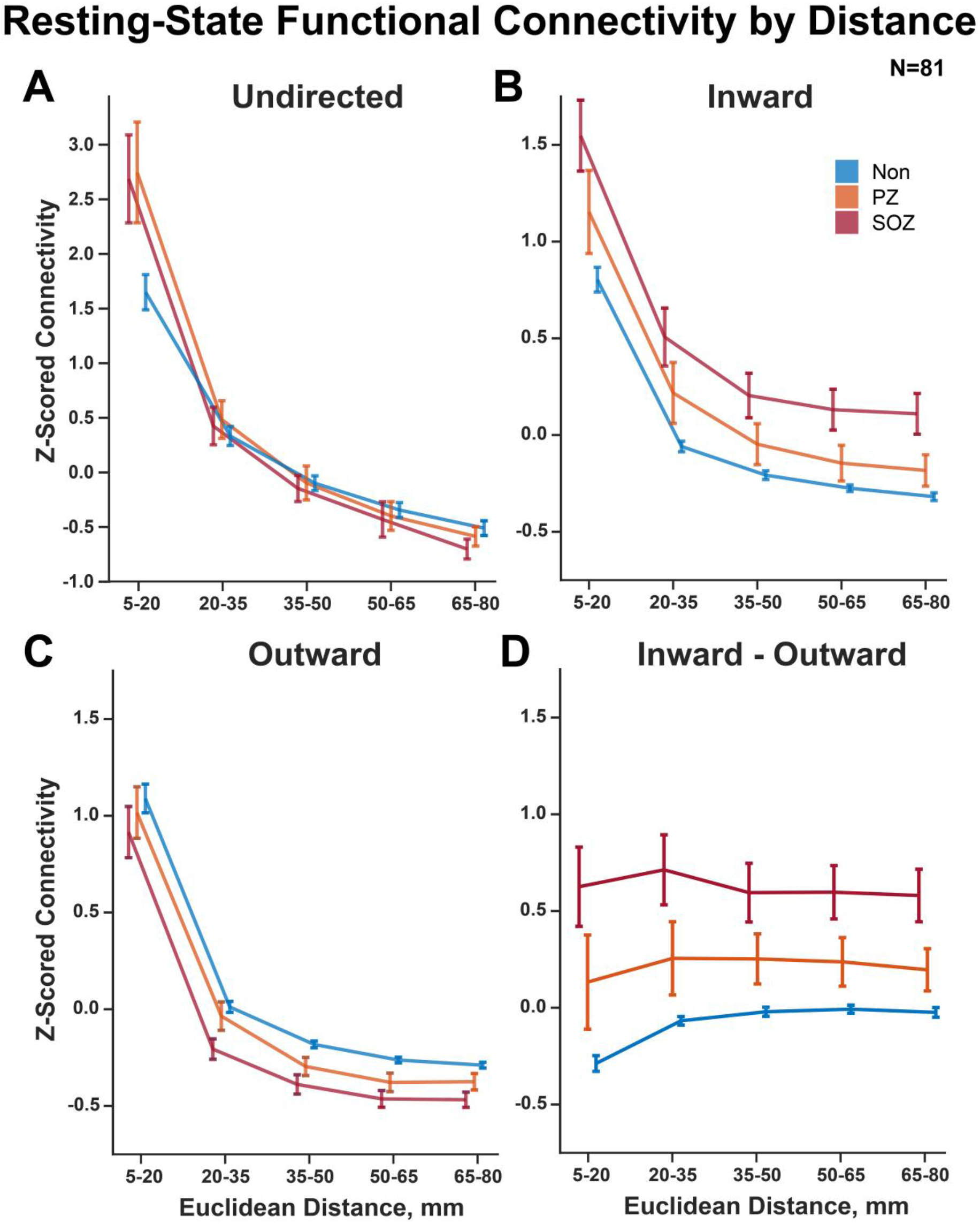
SEEG resting-state connectivity over Euclidean distance. **A-C)** Undirected and directed connectivity declined rapidly with increasing network edge Euclidean distance thresholds (two-way repeated measures ANOVA distance effect p-value<1e-10 for undirected inward and outward connectivity). **D)** Reciprocal connectivity (inward – outward PDC strength) demonstrates a consistent relationship spanning all distance thresholds measured (two-way repeated measures ANOVA group effect p-value=2.6e-12, interaction p-value=1.15e-6). Error bars represent 95% confidence intervals of the mean.

### SOZs Exhibit Local Structure-Function Hypercoupling

Using the SWiNDL technique on 26 patients with DWI available, we observed that SOZs and PZs demonstrated comparably increased structural connectivity compared to non-involved regions despite only SOZs exhibiting significantly increased reciprocal functional connectivity in this cohort (**Figure 4A**, Structural: one-way ANOVA p-value=5.18e-7, SOZ-PZ p-value=1.2e-3, SOZ-Non p-value=3.30e-7; Functional: one-way ANOVA p-value=2.92e-8, SOZ-Non p-value=7.88e-7, PZ-Non p-value=5.13e-7). This suggests that SOZs have enhanced functional suppression to PZs despite a comparable SEEG-specific structural connectivity.

**Figure 4:**
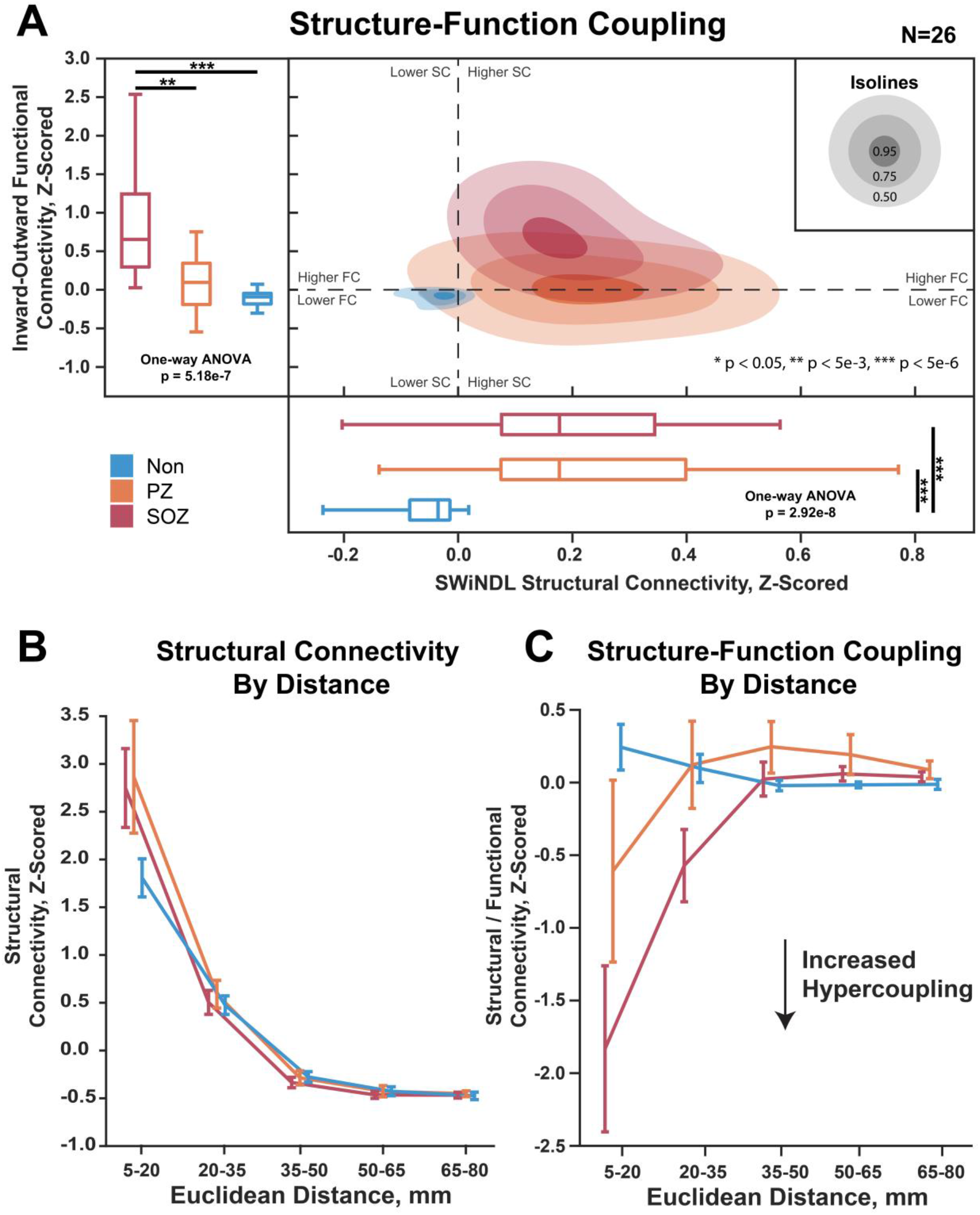
Structure-function coupling over Euclidean distance. A) The left boxplots show the functional connectivity (partial directed coherence inward strength minus outward strength) of seizure onset zones (SOZ), propagative zones (PZ), and non-involved regions (Non). The bottom boxplots show structural connectivity as measured by SWiNDL (Subsampling Whole-brain tractography with intracranial EEG Near-field Dynamic Localization). The contours in the middle show the 2D distribution of the scatterplot of functional vs. structural connectivity. B) Structural connectivity over Euclidean edge distance. Error bars represent 95% confidence intervals. Repeated measures two-way ANOVA: SOZ/PZ/Non effect p=7.52e-3, distance effect p=1.75e-64, interaction effect p=5.44e-9. C) Structural/Functional connectivity over Euclidean edge distance. Repeated measures two-way ANOVA: SOZ/PZ/Non effect p=7.27e-4, distance effect p=8.63e-19, interaction effect p=9.76e-21. Post-ANOVA multiple comparison: **p < 5e-3, ***p< 5e-6. N=26 patients.

We then observed that only the local (5-20 mm) structural connectivity of SOZ and PZs appears to be driving the observed increase in structural connectivity over non-involved regions due to the overlapping of 95% confidence intervals of the mean for all distance thresholds greater than 20 mm (**Figure 4B**). However, the structure-function hypercoupling of SOZs was significantly stronger extending out to 35 mm compared to non-involved regions, with PZs exhibiting an intermediate local (5-20 mm) structure-function hypercoupling (**Figure 4C**, two-way repeated measures ANOVA group effect p-value=7.27e-4, distance effect p-value=8.63e-19, interaction p-value=9.76e-21). These results indicate that SOZs exhibit an inherently increased resting-state structure-function suppressive hypercoupling over short to medium range distances despite a comparable structural connectivity to that of PZs.

### PZ Connectivity Differs Based on Surgical Outcome

When re-evaluating resting-state SEEG functional connectivity over distance in subjects with Engel I vs. Engel II-IV surgical outcomes, we observed that PZs exhibited the largest identifiable difference in connectivity profiles. Specifically, whereas PZs exhibit an intermediate connectivity profile to that of SOZs and non-involved regions in subjects with Engel I outcomes (**Figure 5**, two-way repeated measures ANOVA group effect p-value = 1.16e-2, distance effect -value=1.26e-2, interaction p-value=5.15e-2), PZ 95% confidence intervals of the mean become indistinguishable from non-involved regions and significantly different from SOZs for subjects with Engel II-IV outcomes (**Figure 5B**, two-way repeated measures ANOVA group effect p-value=1.73e-4, distance effect p-value=7.83e-1, interaction p-value=3.01e-3). These results could indicate a fundamental difference of PZ connectivity in subjects with Engel II-IV surgical outcome or could reflect different SEEG sampling and subsequently different ictal event interpretation in these subjects.

**Figure 5:**
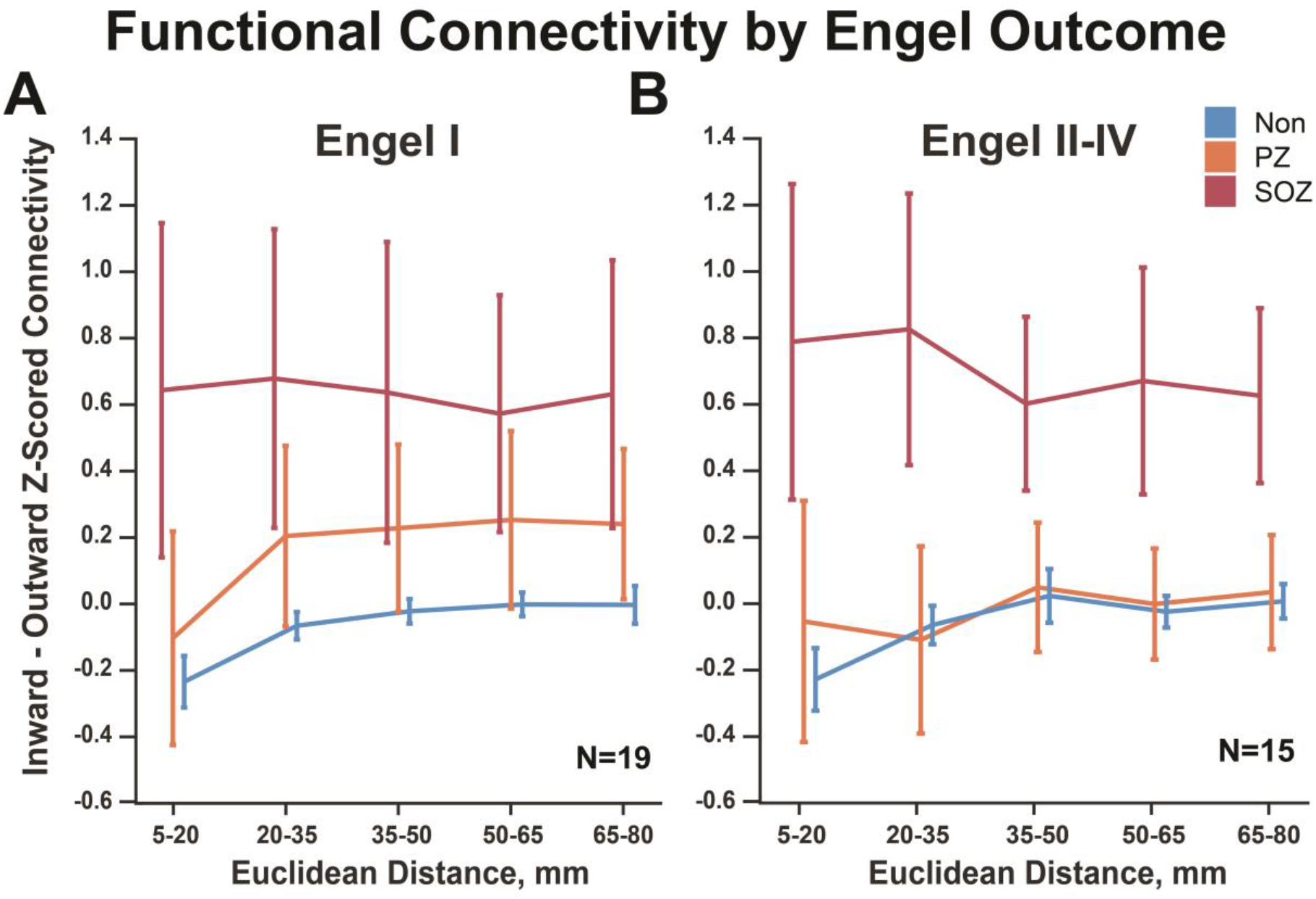
Functional connectivity by Engel outcome. **A)** SOZ, PZ, and non-involved region connectivity for subjects with Engel 1 outcomes (two-way repeated measures ANOVA group effect p-value = 1.16e-2, distance effect -value=1.26e-2, interaction p-value=5.15e-2). **B)** PZs exhibit lower connectivity in subjects with Engel II-IV outcomes (two-way repeated measures ANOVA group effect p-value=1.73e-4, distance effect p-value=7.83e-1, interaction p-value=3.01e-3). Error bars represent 95% confidence intervals of the mean.

### A Structure-Function Coupling Model Most Accurately Classifies SEEG Nodes as SOZ, PZ, or Non-Involved Region

Using a nested cross validation model (**Figure 6A**), we first trained an SVM with only the resting-state SEEG functional connectivity data summarized in **Figure 1** to classify individual SEEG nodes as residing in an SOZ, PZ or non-involved region (**Figure 6B**). The model achieved a completely held out test set overall accuracy of 84.4% (±2.1% standard deviation, SD) and individual true positive rates of 92.0±6.9% for SOZs, 81.6±5.4% for PZs and 53.0±1.8% for non-involved regions. Incorporating SWiNDL structural connectivity derived from DWI data into the model raised the test set overall accuracy to 92.0±2.2% and the individual true positive rates to 92.5±7.2% for SOZs, 93.0±5.3% for PZs and 71.2±2.0% for non-involved regions (**Figure 6C**).

**Figure 6:**
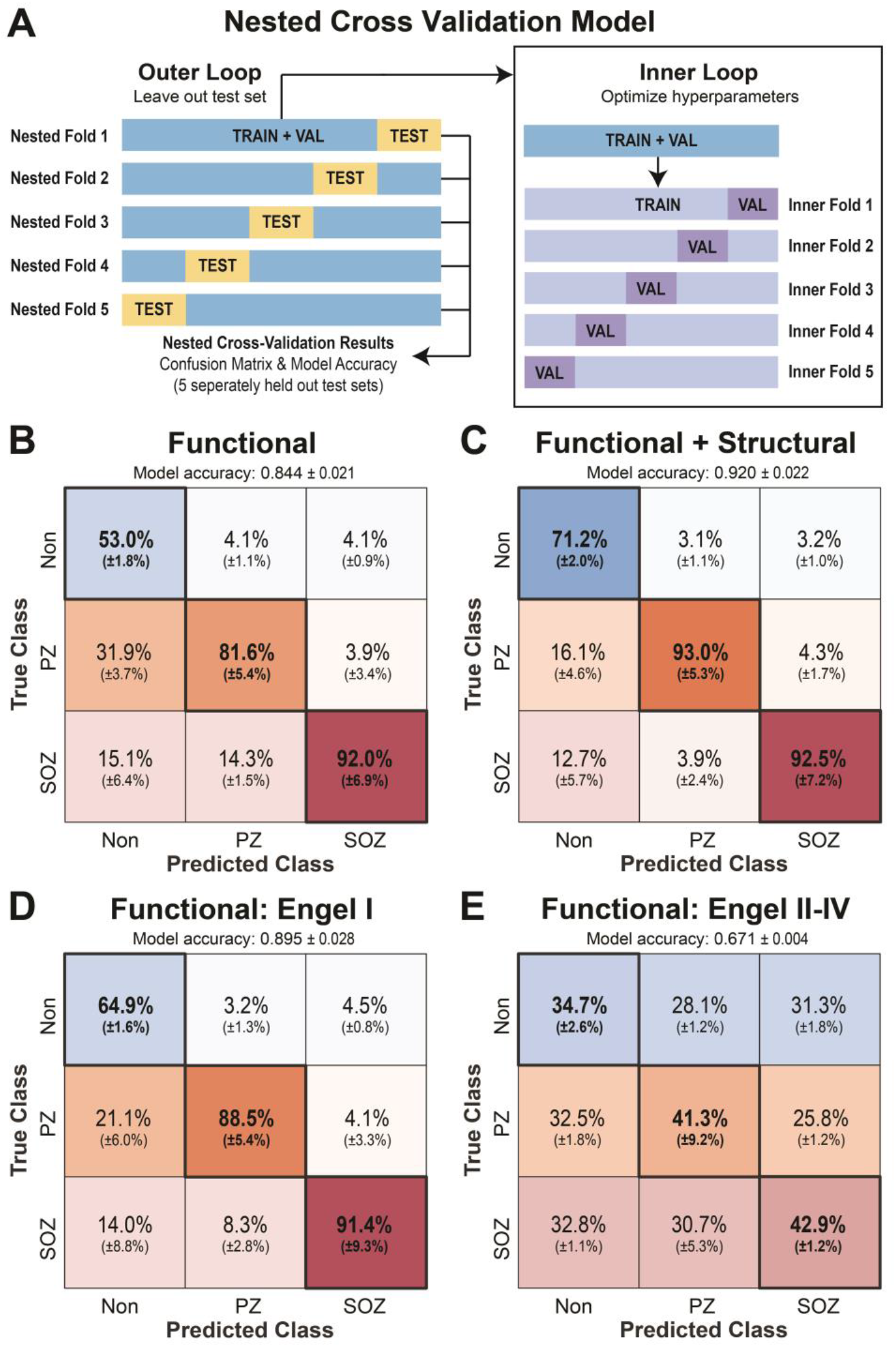
Classification of SOZ, PZ, and non-involved regions using a support vector machine. **A)** A 5-fold nested cross validation scheme was used to evaluate the support vector machine’s ability to classify SOZ vs. PZ vs Non. A completely withheld testing set delineated at the patient level was used for each model evaluation. **B)** Confusion matrix when only functional connectivity was used to generate the model. Overall held-out test set accuracy of 84.4±2.1% (mean ± STD). Confusion matrix percentages are normalized by column – i.e., each confusion matrix entry can be interpreted as “If the model predicts this SEEG contact is a [SOZ/PZ/Non], then there is an X% chance it truly is.” **C)** Confusion matrix for a model using functional and structural connectivity with overall held-out test set accuracy of 0.920±2.2%. **D)** Confusion matrix for a model generated with only Engel I subjects with overall held-out test set accuracy of 89.5±2.8% **E)** Confusion matrix for Engel II-IV subjects tested with the model generated from Engel I subjects (i.e. model “D”) with overall held-out test set accuracy of 67.1±0.4%.

We then sought to test the accuracy of a functional connectivity model only trained on Engel I outcome subjects (**Figure 6D**) – the test set overall model accuracy increased compared to the accuracy for the entire cohort to 89.5±2.8% with true positive rate increasing for PZ and non-involved regions and remaining unchanged for SOZs. Next, when using the Engel I model on Engel II-IV subjects, the accuracies dropped to 67.1±0.4%. Overall, these findings suggest that a model trained on readily available resting-state SEEG data can accurately classify SOZs, PZs, and non-involved regions, but accuracy increases if DWI data are available. Furthermore, the large drop in accuracy when using the Engel I model on Engel II-IV subjects suggests that Engel II-IV subjects have a fundamentally different connectivity profile to that of subjects who achieved Engel I outcomes.

## Discussion

The concept of excitation and inhibition in epilepsy has been investigated for decades.^29, 66–68^ Specifically, at the cellular level, GABA signaling is thought to play a pivotal role in seizure initiation and termination when investigated in vitro, ex vivo, and with optogenetics.^69–71^ When abstracted to electrographic networks, the concept of inhibition and excitation has been explored as it relates to inward and outward connectivity of epileptogenic zones. Recent evidence has suggested that epileptogenic zones have increased inward connectivity which could relate to interictal suppression of epileptiform activity.^19, 20, 72–78^ Whereas other works have focused on observations of increased outward connectivity from SOZs to the rest of the network.^31, 79, 80^ In response to the existing variety of observations and interpretations of functional connectivity in the epileptic network, this work sought to test the specific hypothesis of interictal suppression of SOZs (the ISH) at the electrographic network level by characterizing the direction and relative elevation or attenuation of neural activity.

Properly testing the ISH with only resting-state SEEG connectivity is challenging due to the difficulty of interpreting whether resting-state functional connectivity is excitatory, inhibitory, or neutral in nature. For example, increased inward connectivity could be interpreted with known directed connectivity motifs,^81^ but it will not directly indicate if this increase in inward connectivity correlates to any region being suppressed or excited. Thus, we conducted SPES to 1) provide additional insight into the directionality of network connectivity with another paradigm, and 2) quantify elevation or attenuation of frequency band power by measuring the change in relative band power from pre-stimulation baseline. Using SPES, we can gain insight into the potential excitatory and inhibitory nature of the network connectivity. However, it is important to consider that SPES itself may be altering the network dynamics away from the resting state. Thus, interpretation of resting-state SEEG and SPES findings must be carefully integrated.

Further complicating this discussion, one cannot assume that attenuation in a certain frequency band power aligns with inhibition. As an example, there is a large body of evidence that an increase in beta power correlates to inhibition of motor circuits, particularly in Parkinson’s disease.^82, 83^ For the sake of clarity, we will use the term ‘elevation’ to refer to an increase in signal band power, and ‘attenuation’ to refer to a decrease. We will reserve the terms ‘excitation’ and ‘inhibition/suppression’ for the concept of how a modification in neural activity relates to a change in a clinically observable behavior of interest (e.g. seizing or not seizing). Furthermore, we use the term “seizure onset zone” to indicate an area of first observed electrographic epileptiform changes. We reserve the use of the term “epileptogenic zone” (i.e. “site of the beginning of the epileptic seizures and of their primary organization”) for discussion of a hypothetical electro-clinically defined region that could be resected to render the subject seizure free.^10^ With these considerations and definitions in mind, we will discuss our interpretation of the pertinent findings in this work.

### Does Imbalanced Reciprocal Connectivity Indicate Suppression?

In this study, we observed imbalanced reciprocal (inward – outward) connectivity of SOZs and PZs: Specifically, SOZs demonstrated a large increase in inward connectivity and a moderate decrease in outward connectivity for resting-state SEEG analyses. PZs exhibited a similar imbalance in reciprocal connectivity to SOZs, but to an intermediate extent. Important for interpretation, non-involved region reciprocal connectivity was approximately zero – i.e. regions not directly involved in observed seizure onset or early propagation have roughly equal inward and outward connectivity. This suggests that SOZs and PZs interact with the network differently than presumably healthy regions of the brain. One interpretation of this finding is that SOZs are actively isolated/segregated by other regions of the brain. It is possible that healthy regions are signaling for the SOZs and PZs to be inhibited (increased inward connectivity) and thus suppress their ability to communicate with the rest of the network and initiate seizures (decreased outward connectivity).

Further evidence for SOZ segregation was exhibited in the reciprocal connectivity elucidated with low-frequency stimulation. Using a power spectral density approach,^74, 84, 85^ we intended to build on previous works that focused on whole-band power metrics like root mean square,^77, 79, 86^ or metrics that rely on cortico-cortical evoked potential waveform morphology which can be difficult to interpret with depth electrodes.^75, 87–91^ Overall, when presumably healthy regions were stimulated, the SOZ theta [4-8 Hz] relative band power was markedly decreased from pre-stimulation baseline. Conversely, SOZ beta [13-30 Hz] and gamma [31-80 Hz] relative band power increased when healthy regions were stimulated. Increased inward evoked gamma band power in SOZs is in alignment with previous low-frequency stimulation findings.^74^ Notably, reciprocal evoked alpha band power was not altered across all region designations, and PZ relative band power was not significantly altered for any frequency band measured.

It is long known that theta spectral power accounts for a significantly larger portion of whole band power on EEG recordings compared to beta and gamma power.^92^ Additionally, theta power is thought to be involved with long range integration of brain regions, whereas gamma power is considered important for local integration.^93–95^ Thus, simultaneous attenuation of a high-power spectral band (theta), that is also implicated in long range integration, in parallel with excitation of a locally integrating frequency band (gamma), suggests that SOZ functional segregation is increased when non-involved regions are stimulated. Furthermore, gamma activity is thought to reflect GABA inhibitory interneuron activity.^96–98^ Thus, increased inwardly activated gamma band power in SOZs could reflect a direct change in the GABA interneuron balance. Additionally important to consider is that Gamma activity, in the form of low-voltage fast activity (LVFA), is commonly observed on intracranial EEG immediately before ictal onset and increased baseline gamma power is thought to be a biomarker for the “ictal core”.^99–101^ Thus, it is possible that the observed SOZ increase in gamma power when non-SOZ nodes are stimulated is simply SOZs demonstrating an enhancement of an intrinsic pathologically distinct phenomenon and could actually be driving SOZs toward an ictal state. Regardless, the low-frequency stimulation findings provide corroborating evidence for an electrographic network-level increase in inward connectivity to the SOZ and provide insight into the frequency-specific behavior of SOZs during stimulation.

### Reciprocal Connectivity is Edge Distance Invariant, but SOZs Exhibit Local Structure-Function Hypercoupling

We sought to investigate if findings of increased inward connectivity to SOZs were affected by Euclidean distance and/or estimates of anatomical white matter connectivity between SEEG contacts. Past work has demonstrated the importance of considering Euclidean edge distance when analyzing functional epileptogenic networks. Specifically, it has been demonstrated that short-range structural connections drive the majority of aberrant SOZ connectivity and could dictate seizure spread.^19, 102^ This observation of rapid functional connectivity decay with increasing edge length was recapitulated in this work’ s undirected and directed resting-state findings. However, long range connections are thought to add diversity and complexity to brain networks and could be important contributors to widescale integration.^103^ Concordantly, our finding that resting-state reciprocal connectivity is relatively constant across distance suggests that distant brain regions may still play an important role in SOZ excitation and inhibition.

Beyond simple Euclidean distance metrics, diffusion-derived structural connectivity characteristics can be used to investigate potential pathological SOZ connectivity.^104–110^ We observed SOZs and PZs to have increased structural connectivity relative to non-involved regions, which is in alignment with past SEEG-informed diffusion imaging studies.^105^ This increase could represent an intrinsic pathological biomarker of the epileptogenic network or could possibly be due to implantation bias increasing the density of SEEG contacts around suspected SOZs. Of more pertinent interest to testing the ISH, we investigated if structural connectivity variation could explain our observed differences in functional connectivity. The “coupling” between structure and function is the term commonly used when the strength of structural connections predicts the strength of functional connections.^111^ Thus, a region exhibits strong structure-function coupling when the ratio of their relative functional and structural connections is unity. We use the term “hypercoupling” to refer to connections that exceed the functional connection strength expected from structural connectivity. We observed that SOZs demonstrate markedly increased local hypercoupling – i.e. the increased inward functional connectivity strength is disproportionally elevated above the average structure-function coupling by two standard deviations for edge connections within 5-20 mm. This suggests an abnormal functional reorganization around the SOZs disproportionate to any alterations in structural connectivity.

### Proper PZ Identification May Help Prognosticate Surgical Outcome

A sub analysis with Engel I vs. Engel II-IV surgical resection outcomes demonstrated that PZ connectivity resembled non-involved region connectivity in Engel II-IV subjects. This could be due to an intrinsic difference in PZ connectivity between surgical responders and non-responders, a difference in SEEG implantation strategy between the patient groups that affects network observations, or a difference in ictal interpretation that affects node designation.^112–114^ With this observation as motivation, we developed SVM models to classify SOZs, PZs, and non-involved regions. The models were able to classify SOZs with very high accuracy but differed in ability to differentiate PZs from non-involved regions. Notably, our SVM model to classify individual SEEG bipolar pairs confused PZs and non-involved regions when the entire subject cohort was used in model generation and testing. When restricted to Engel I subjects, the model significantly improved classification between PZs and non-involved regions, with subsequently very poor performance when this model was used to test Engel II-IV subjects. Of note, the best model was produced when diffusion-derived metrics were included, but these data are less commonly collected during presurgical workup. Overall, the models very accurately classified SOZs, but we encourage other groups to include PZ classification into model design because it may be an important factor for prognosticating surgical outcome.

### Limitations

This work utilizes multiple analysis techniques to test the ISH but is limited by the ambiguity of network neuroscience interpretation and integration. Electrographic phenomena observed on SEEG and probabilistic tractography performed on diffusion imaging are removed from underlying neurophysiology and neuropathology. Thus, interpretation of functional and structural network connectivity findings must resist the proclivity toward self-justified and circular conclusions. Further complicating interpretation is the lack of control groups for these populations. SEEG control data does not exist, and we did not include neuroimaging controls. All scientific inferences and hypothesis testing were conducted on comparisons to regions presumably not involved in the genesis or early propagation of epileptiform activity. For subjects with epilepsy, it is reasonable to assume that the entire brain network is sick and thus intra-patient normalization to “healthy” regions could be an ill-posed analytical model. As more depth electrodes are being used for therapy in people with varying psychiatric and neurological disorders, large-scale data sharing to expand our understanding of baseline electrographic phenomena across a more diverse population could help address these issues.

### Conclusions

In summary, we observed electrographic evidence to support the interictal functional suppression of SOZs and PZs on resting-state SEEG analyses and low-frequency stimulation. When controlling for distance and structural connectivity we found that SOZs and PZs demonstrate consistent long-range relative reciprocal connectivity, but elevated local hypercoupling of functional connectivity that could indicate enhanced interictal suppression of SOZs. Finally, we have demonstrated that a clinically useful model can be generated using only SEEG data to classify SOZs, PZs, and non-involved regions. However, inclusion of DWI-derived structural connectivity can increase the model accuracy. Overall, we believe this work supports the hypothesis of interictal suppression of SOZs and could have practical implications to reduce the morbidity of the presurgical workup and aid clinical decision making for this devastating neurological disorder.

## Notes

### Competing Interest Statement

The authors have declared no competing interest.

### Summary of Updates

license update

